# Integrating experiments and simulations to unravel coacervate-membrane Interactions: Insights into de-mixing and morphology modulation

**DOI:** 10.1101/2025.05.16.654554

**Authors:** Sayantan Mondal, Agustín Mangiarotti, Rumiana Dimova, Qiang Cui

## Abstract

Intrinsically disordered proteins and polypeptides can undergo liquid-liquid phase separation (LLPS) to form condensates/coacervates, which play numerous regulatory roles in the cell. Recently, the relevance of such LLPS occurring in the vicinity of membranes has been brought to light by several experimental studies. Membrane-adsorbed condensates are crucial for biomolecular localization, and in some cases, phase separation of proteins at the membrane surface induces significant changes in membrane morphology. A detailed microscopic understanding of the mechanisms behind these observations remains incomplete. Here we combine experiments and molecular simulations to unravel structural and dynamic features of the coacervate/membrane interface across scales. We study poly-Lysine/poly-Aspartate (K_10_/D_10_) coacervates as a prototype of phase-separated condensates with different unilamellar liposomes. Using a multiscale characterization approach that combines confocal microscopy, hyperspectral imaging, fluorescence recovery after photobleaching, and two complementary coarse-grained approaches, we show that the membrane-condensate affinity can be tuned by the anionic lipid content and quantified through the intrinsic contact angle – a material property derived from system geometry – both *in vitro* and *in silico*. We find that the membrane region in contact with the condensate displays a nearly two-fold reduced fluidity compared to the bare membrane. This is attributed to orientational ordering of lipid tails, resulting in decreased area per lipid. Moreover, we observed local lipid de-mixing induced by the coacervate adsorption. This study provides an effective framework for integrating experiment and computation to characterize the properties of coacervate/membrane interfaces that are critical to the functional impacts of these interactions.

## INTRODUCTION

The liquid-liquid phase separation (LLPS) of proteins and genetic material leads to the formation of biomolecular condensates.(1) These membraneless organelles (such as nucleolus, stress granules, processing bodies, etc.) provide an additional means to compartmentalize subcellular processes(2) and are involved in diverse functions, ranging from genome reorganization and transcription(3–5) to stress response(6, 7) and signal transduction.(8) During the past decade, there has been an increasing interest in comprehending the formation, internal structure, and dynamical behavior of bio-molecular condensates, both experimentally and theoretically.(9–21)

The intrinsically disordered regions (IDRs) in proteins and RNAs have been identified as the participants and key regulators of LLPS in biological systems. They can establish multivalent interactions, either homotypic or heterotypic, leading to LLPS.(22, 23) In this context, complex coacervates of polyelectrolytes/polyampholytes assembled by associative phase separation are used as models of phase-separated droplets because they capture key features of membraneless organelles and allow the development of biomimetic systems for synthetic biology applications.(24–27)

In recent years, several biological processes involving the interaction of biomolecular condensates and membrane-bound organelles have been described.(28–32) This phenomenon, known as wetting,(33–35) can promote the mutual remodeling of membranes and condensates,(36–42) leading to a variety of elastocapillary phenomena.(43, 44) In this regard, experiments *in vitro* and *in silico* with model systems permit the precise control and tuning of the physicochemical properties of both membranes and condensates allowing for visualization and quantification of the resulting morphologies. When in contact with giant unilamellar vesicles,(45, 46) lipid-anchored proteins can undergo two-dimensional phase separation at the membrane surface, leading to inward tubulation in vesicles due to compressive forces.(38) On the other side, non-anchored condensates undergo wetting transitions upon contact with membranes, resulting in mutual remodeling of the membrane and condensate.(33, 36, 47, 48) In the presence of excess membrane area, the condensate-membrane interface can become ruffled, forming finger-like protrusions, indicating that condensates can promote complex membrane remodeling.(48) In addition, recent studies have shown that membrane wetting by biomolecular condensates can facilitate the endocytosis of the droplets,(49–52) as well as influence the kinetic and thermodynamic coupling of the lipid and condensate phases.(53, 54) This interaction also influences lipid packing and hydration, even to the extent of inducing phase separation in the membrane.(48, 50, 55)

At the molecular scale, contact with a membrane can influence the conformational ensemble of IDPs, potentially altering their phase behavior. The single-chain properties (such as θ-temperature and Boyle temperature) of an IDP were shown to be correlated with the condensate properties (such as the critical temperature).(56) For example, fused in sarcoma (FUS) forms fibrillar β-sheet rich structures on phosphatidylserine membranes, whereas it forms entangled condensate on a phosphatidylglycerol membrane.(47, 57) Hence, predicting IDP phase behavior upon membrane wetting cannot be solely based on increased local concentration. Furthermore, the phase behavior of membranes and IDPs can be coupled.(53, 58, 59) Membranes can promote LLPS at their surface, even under conditions unfavorable for LLPS in bulk solutions.(60, 61) These studies highlight the importance of membrane surfaces as critical regulators of LLPS and condensate properties.(39, 62) Furthermore, LLPS provide a novel mechanism for membrane remodeling in cells, complementing established mechanisms such as hydrophobic insertion, scaffolding, protein crowding, and coating.(63–65)

Interaction between polyelectrolyte/polyampholyte coacervates and membranes have been shown to lead to significant membrane remodeling. For example, polyelectrolyte coacervates made of an equimolecular mixture of E_30_ and K_30_ can induce a pronounced negative curvature and local lipid demixing when partially wetting an anionic lipid bilayer.(36) Further studies demonstrate that the extent of such membrane remodeling depends on the charge patterning of the IDR chains.(40) Continuum mechanics-based descriptions suggest that the increased inter-protein contacts, driven by negative curvature, are the primary drivers of the observed membrane remodeling.(38) However, recent evidence demonstrates that such hypothesis may not hold in case of coacervate-membrane interaction where counterions play an important role in providing enthalpic stabilization.(36)

Although several studies have explored specific aspects of the interaction of polyelectrolyte coacervates with membranes using either *in vitro* or *in silico* approaches,(36, 37, 49, 50, 66) a systematic and integrative investigation exploiting the strengths of both methodologies is lacking. Here, we bridge this gap by combining experiments with molecular dynamics, providing a multiscale approach to unravel the effects of coacervate wetting on membrane properties and remodeling.

## MATERIALS AND METHODS

### A. EXPERIMENTAL PROCEDURES

#### Materials

The phospholipids 1,2-dioleoyl-sn-glycero-3-phosphocholine (DOPC) and 1,2-dioleoyl-sn-glycero-3-phospho-L-serine (DOPS) were purchased from Avanti Polar Lipids (IL, USA). ATTO 647N-DOPE was obtained from ATTO-TEC GmbH (Siegen, Germany). Chloroform HPLC grade (99.8 %) was purchased from Merck (Darmstadt, Germany). All lipid stocks were prepared in chloroform at 4 mM, containing 0.1 mol% ATTO 647N-DOPE, and stored until use at-20°C. Sodium chloride (NaCl), potassium chloride (KCl) and magnesium chloride (MgCl_2_) were obtained from Sigma-Aldrich (Missouri, USA).

The oligopeptides poly(l-lysine hydrochloride) (K_10_) and poly(l-aspartic acid sodium salt) (D_10_), each with a degree of polymerization n=10, were purchased from Alamanda Polymers (AL, USA) and used without further purification (purity ≥ 95%). The N-terminal TAMRA-K10 was purchased from Biomatik (Ontario, Canada). All solutions were prepared using ultrapure water from SG water purification system (Ultrapure Integra UV plus, SG Wasseraufbereitung) with a resistivity of 18.2 MΩ cm.

#### Condensate formation

Phase separation was induced by mixing equimolar aliquots of K10 and D10 (5 mM) in a solution containing 15 mM KCl, 0.5 mM MgCl_2_, and 170 mM glucose. For labeling, TAMRA-K_10_ was added at a concentration of 1 mol%. The final osmolarity of the mixture was ≈ 200 mOsm which was adjusted using a freezing point osmometer (Osmomat 3000, Gonotec).

#### Vesicle preparation

GUVs were grown using the electroformation method. Briefly, 3-4 µL of the lipid stock were spread on two conductive indium tin oxide-coated glasses and kept under vacuum for 1 h. A chamber was formed using a rectangular 2-mm-thick Teflon frame sandwiched between the two glass electrodes. The lipid films were hydrated with 2 ml of an isotonic sucrose solution (matching the osmolarity of the condensate solution). An electric AC field (1V, 10 Hz, sinusoidal wave) was applied for one hour at room temperature. Once formed, the suspension of the giant vesicles was stored at room temperature until use. The vesicles were prepared freshly before each experiment.

#### Condensate-membrane suspensions

For the interaction of membranes with K_10_/D_10_ condensates, the vesicle suspension was diluted 1:10 into the final buffer matching the droplets suspension conditions. Then, an aliquot of this solution was mixed with the droplet suspension at an 8:1 volume ratio directly on a coverslip. The sample was then sealed for immediate observation under the microscope.

#### Confocal microscopy and FRAP

A Leica SP8 confocal microscope equipped with a 63x, 1.2 NA water immersion objective (Mannheim, Germany) was used for imaging. TAMRA-K10 and ATTO 647N-DOPE were excited using the 561 nm and 633 nm laser lines, respectively. For FRAP measurements, a circular region of interest (ROI) with a diameter of 2 μm was elected at the vesicle equator and photobleached during 3 iterative pulses over a total duration of ∼3 s. The fluorescence recovery within the ROI was recorded and analyzed using ImageJ.

#### Contact angle determination

Confocal projections enable the determination of three apparent microscopic contact angles at the contact line between the vesicle and the condensate droplet, *θ*_*e*_ (opening toward the external solution, *θ*_*c*_ (toward the condensate phase), and *θ*_*vv*_ (toward the vesicle interior). Accurate measurements of these contact angles between the different surfaces of the bare and wetted membrane segments and the condensate interface, require that the rotational axis of symmetry of the vesicle-droplet system lies in the imaging plane. This alignment is crucial to ensure precise geometry and contact angle determination, as explained in detail in Ref (37). Misalignment can lead to incorrect assessment of the system geometry and contact angles.

The geometric factor characterizing the membrane wetting affinity is defined as(37, 67,68)

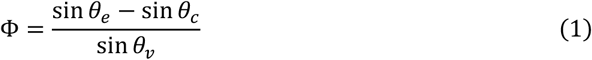

Φ can vary from +1 (a de-wetted state) to −1 (complete wetting). At the nanometer scale, the membrane is smoothly curved,(69) and the wetting is characterized by the intrinsic contact angle (*θ*_*in*_) (70), see below.

#### Spectral phasor analysis

Hyperspectral images of vesicles labeled with LAURDAN were analyzed using the spectral phasor method.(71) This analysis calculates the real (G) and the imaginary (S) component of the Fourier transform. The Cartesian coordinates (G and S) of the spectral phasor plot are defined by the following expressions

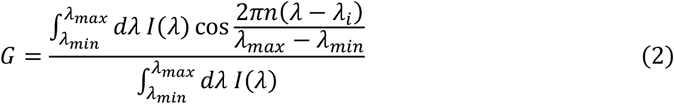

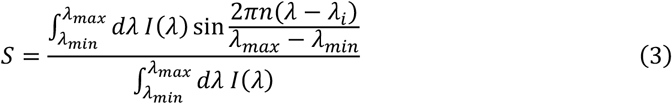

where, for a given pixel, I(λ) is the pixel intensity at wavelength λ, *n* denotes the harmonic number, and λ_i_ is the initial wavelength.

The spectral phasor approach follows the rules of vector algebra, known as the linear combination of phasors. This property implies that a combination of two independent fluorescent species (or states) will appear on the phasor plot at a position that is a linear combination of the phasor positions of the two independent spectral species. In this manner, two-cursor analysis was used to calculate the histogram for the pixel distribution along the trajectory for changes of LAURDAN fluorescence. The histograms represent the number of pixels at each step along the line between two cursors, normalized by the total number of pixels. We plotted the average value for each histogram ± standard deviation, as well as the center of mass of the histogram for quantitative analysis with descriptive statistics. The center of mass was calculated following:

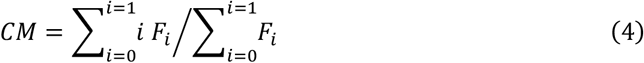

where F_i_ is the fluidity fraction.

#### Data reproducibility and statistics

At least three independent experiments were used to perform statistical analysis. Histograms are shown as means ± standard deviation (SD). The center of mass is represented as scatter plots containing the individual measurements and the mean values (± SD). Results were analyzed using one-way ANOVA with Tukey post-test, where p < 0.05 was considered statistically significant. Statistical analyses and data processing were performed with the Origin Pro software (OriginLab corporation).

### B. HEORETICAL AND COMPUTATIONAL PROCEDURES

#### Computational modeling

Because of the vast conformational space of IDRs, molecular dynamics (MD) simulations in the atomistic resolution (with explicit molecular water and ions) are computationally expensive. Therefore, one often resorts to different levels of coarse-grained (CG) approaches,(72, 73) for example, single-bead per amino-acid residue, multiple-beads per residue, single-bead for multiple residues, etc. Here, we have used CG modeling of two different resolutions, namely, (i) a generic model that consists of Cooke three-bead lipids(74) with bead-spring polymers; and (ii) MARTINI 3 model with explicit water/ion beads.(75) The primary caveat of CG approaches is the loss of atomistic and dynamic information. However, CG models can efficiently provide structural information at biologically relevant length-scale and long timescale that are otherwise inaccessible to atomistic MD.

#### The generic coarse-grained model

The generic model consists of a liposome made of three-bead Cooke lipids (**Figure 1e**, *right panel*), and two types of polymer chains (A_10_ and B_10_) in equimolecular proportions that represent the polyelectrolytes (K_10_/D_10_). This setup has no explicit beads for water and ions. Such models have been successful in recapitulating several experimentally observed phenomena such as coacervate engulfment and multilayered membrane formation.(50) The polymers are modeled as self-repulsive chains implemented through the Weeks-Chandler-Anderson (WCA) potential(76) with heterotypic attraction implemented through the Lennard-Jones (LJ) potential described by Eq.(5) and Eq.(6), respectively

**Figure 1.**
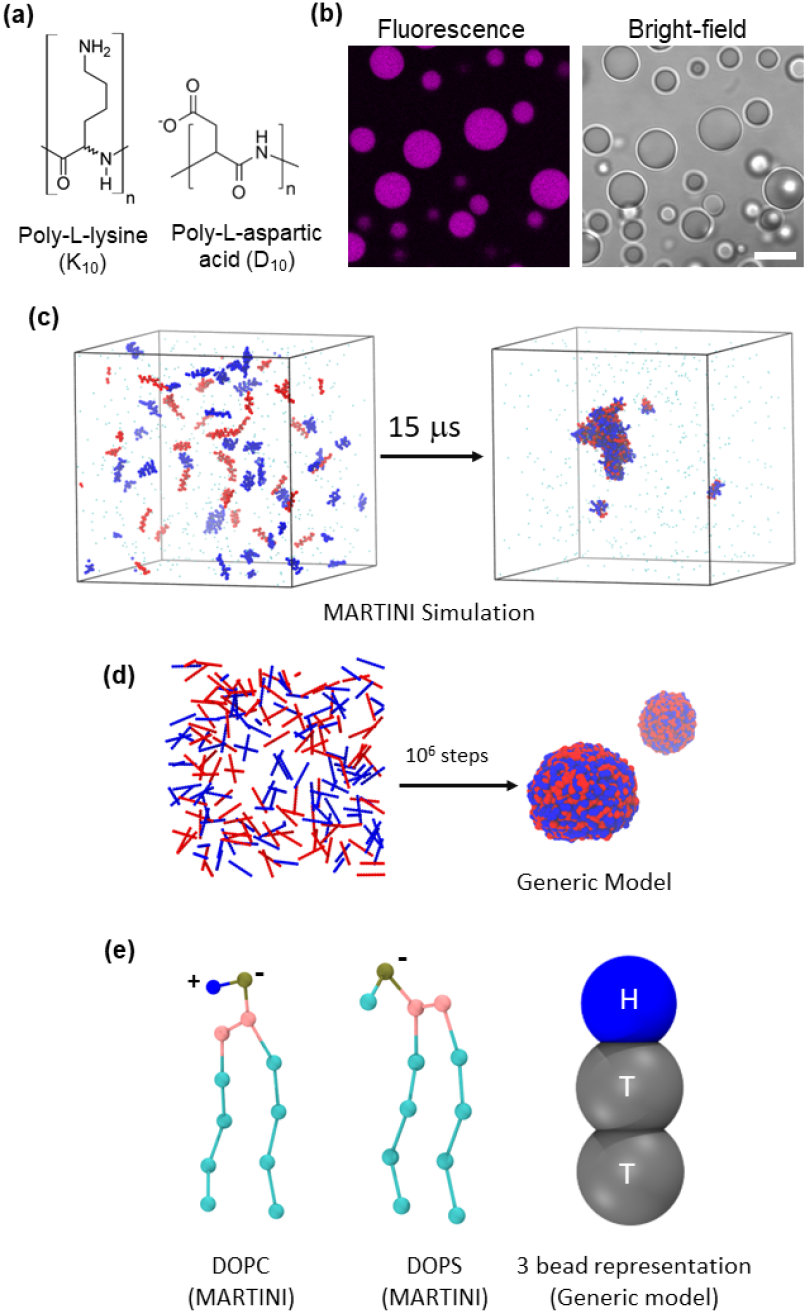
Molecular structures, models and coacervate formation in experiments and simulations. (a) Molecular structures of the poly-lysine (K_10_) and poly-aspartic acid (D_10_) oligopeptides. The number of monomers (n) is 10. (b) Confocal fluorescence microscopy and bright-field images of coacervates formed by the equimolar mixture of K_10_ and D_10_ peptides (∼5 mM) in 15 mM KCl and 0.5mM MgCl_2_. A 1 mol% of labeled peptide (K_10_-TAMRA) was included for visualization [Scale bar=20 µm]. (c) Coacervation of an equimolecular mixture of K_10_ and D_10_, as observed in MARTINI based coarse-grained simulation for 15 μs. (d) Coacervation observed from a generic CG model where the amino acids are represented by single beads (‘A’ and ‘B’) which are self-repulsive (A-A and B-B) but heterotypically attractive (A-B). (e) Lipid models used in this study: MARTINI lipids and the three-bead representation (H-T-T) for the generic CG model.

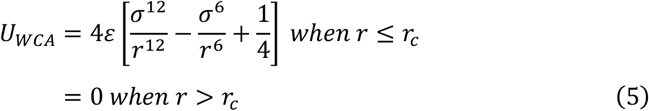

and

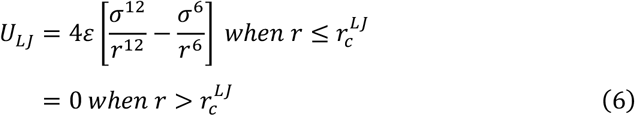

where *r* is the inter-bead distance, *ε* is the well-depth, *r*_*c*_ = 2^1/6^σ and 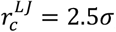, and *σ* is the bead diameter. There are harmonic bonds between successive beads in the same polymer with a dimensionless force constant of 50.0 and the equilibrium bond length *r*_*eq*_ = 0.548. The well depth *ε* is set to be 1.5 between the polymer beads. *σ*_*AA*_ and *σ*_*BB*_ both are set to 0.5. All values for this model are in the reduced unit where 1**σ** ≈ *6*.*93* Å.

Each lipid molecule consists of three beads, one for the head (H) and two for the tail (T). The heads are self-repulsive with WCA potential [Eq. (5)] and the tail beads are attractive according to a combination of truncated LJ and cosine potential, given by:

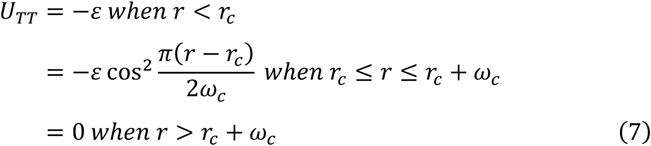

Here, *σ*_*HH*_ = 0.95 = *σ*_*HT*_ and *σ*_*TT*_ = 1.00 (recall that 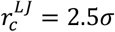). The *ω*_*c*_ parameter for TT, *ω*_*TT*_, could be different depending on the lipid composition and stiffness of the bilayer. The successive lipid beads are linked by finite extensible nonlinear elastic (FENE)(77) bonds [Eq. (8)] with *k* = 30.0 and *r*_∞_ = 1.5

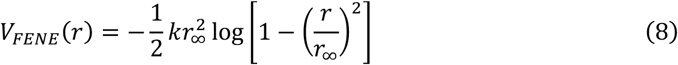

The polymer-head (A-H and B-H) interactions are attractive LJ type, and the polymer-tail (A-T and B-T) interactions are repulsive WCA type. Therefore, we have two tunable parameters to describe different systems with the same model in a mean-field manner, namely, the polymer-head interaction strength (*ε*_*PL*_) and the lipid inter-tail interaction range (*ω*_*TT*_). The rationale behind the choice of the two tunable parameters is the following. When the lipid composition in the vesicle changes, the stiffness (or, bending modulus) of the bilayer as well as the effective attraction between the lipid and polymers will change.

#### MARTINI coarse-grained model

As mentioned above, we model the system using another general-purpose, explicit-solvent, and popular CG force-field, MARTINI (v3.0).(75) MARTINI uses a 4-to-1 mapping strategy where 4 non-hydrogen atoms are represented by one bead. MARTINI-3 was used previously to study the salt-concentration dependence of polyelectrolyte coacervation(78) and later to study coacervate-membrane interaction.(36) In the context of IDPs, a rebalancing strategy has been proposed by scaling the protein-water interactions in MARTINI-3 that improves the single-chain behavior like radius of gyration and end-to-end distance.(79) In addition, the MARTINI model is known to provide a reasonable description of the lipid bilayers.(80) In MARTINI, each D and K residue is modeled by two CG beads, one for the backbone and one for the sidechain. The sidechain bead of each D bears a −1e charge and the side-chain bead of each K bears a +1e charge (e=charge of a proton). The N-and C-termini, respectively, bear +1e and −1e charges. For the lipid bilayer, we have used two different kinds of lipids, namely, DOPC and DOPS, in different ratios. The details of parameterization can be found in the original MARTINI 3 paper.(75)

#### Molecular dynamics simulation details

For the generic CG model, we first simulate 1000 copies of each polymer chain inside a (40 unit)^3^ cubic box to form the droplet (**Figure 1d**). We separately prepare a unilamellar vesicle of 50 unit diameter with approximately 9500 three-bead lipid molecules inside a (70 unit)^3^ cubic box. After that, the preformed droplet is placed on the outside of the vesicle. The systems are first energy minimized and then equilibrated for 10^5^ steps in an NVT ensemble with a timestep (Δt*) of 0.002 (in reduced units), during which the initial adsorption occurred. Following this, we perform production runs for another 10^7^ steps in an NVT ensemble with Δt*=0.005. We use a Langevin thermostat to keep the average reduced temperature (T*) at 1.1 with a damping timescale of 50.0. The coordinates are recorded in every 10^4^ steps. For these simulations, we used the LAMMPS(81) package and VMD(82) for visualization.

For the MARTINI simulations, the atomistic versions of the polyelectrolytes (K_10_ and D_10_) are prepared using *pymol*,(83) followed by the conversion in MARTINI-CG format using the *martinize2* script.(84) Mimicking the experimental conditions, we first simulate 33 copies of each polyelectrolyte inside a (30 nm)^3^ cubic box filled with 226,627 MARTINI water beads, 271 Na^+^ and 271 Cl^-^ ion beads. This is done for 15 μs to check the LLPS propensity in the bulk (**Figure 1c**). Then we place the preformed droplet near different equilibrated lipid bilayer patches of dimension (32 nm)^2^. However, the original MARTINI force field was unable to capture the experimentally observed adsorption of the coacervate on the DOPC bilayer and showed weak/transient adsorption with the 20% DOPS bilayer, suggesting that the interactions between polyelectrolytes and lipids are underestimated. Therefore, we re-scale some of the force field parameters by scaling up the interactions between the polyelectrolytes and lipid heads. A similar technique was adapted earlier in the context of the membrane protein association.(85) Here we scale the non-bonded interaction strength between the protein beads and the lipid headgroup beads (four of them) of DOPC lipids by a factor of 1.20. On the other hand, due to the presence of Coulombic attraction between DOPS and polyelectrolytes, we do not scale the LJ parameters associated with DOPS charged head. We first minimize the energy of the systems followed by 100 ns equilibration in an NpT ensemble (T=298 K and p=1 bar). After that a 5 μs production run is carried out with the same T and p. All equilibration simulations are propagated with a time step of 10 fs and production simulations are propagated with a time step of 20 fs, using the leap-frog algorithm. We use the V-rescale thermostat (τ_T_ = 1 ps^−1^) at 298 K and Parrinello–Rahman barostat with semi-isotropic pressure coupling (τ_P_ = 12 ps^−1^) at 1 bar to make the bilayer tensionless. For initial equilibration purpose, we use Berendsen barostat with τ_P_ = 6 ps^−1^. The electrostatic interactions are screened with a reaction-field (ε_r_) of 15 within a cut-off of 1.1 nm and non-bonded interactions are also terminated at 1.1 nm with the Verlet cut-off scheme. The coordinate dumping rate is set to 100 ps. For all the MARTINI simulations, we use the GROMACS 2018.3 simulation package(86) and VMD(82) for visualization.

## RESULTS AND DISCUSSIONS

### 1. Membrane wetting and remodeling by peptide coacervates

We formed coacervates of the oligopeptides poly-L-lysine (K_10_) and poly-L-aspartate (D_10_) (see **Figure 1a**) in a solution containing 15 mM KCl and 0.5 mM MgCl_2_, as previously described.(24, 37) **Figure 1a** shows the molecular structure of the polypeptides. By using a 1 mol% of TAMRA-K10, the coacervates can be observed with fluorescence confocal microscopy, as shown in **Figure 1b**. In the simulations, the polyelectrolyte mixtures underwent LLPS with both MARTINI-3 (**Figure 1c**) and the generic model (**Figure 1d**) force fields.

To evaluate the coacervate interaction with membranes we used giant unilamellar vesicles (GUVs) composed of pure DOPC and DOPC:DOPS mixtures, in the presence and absence of sodium chloride (NaCl). For observation by confocal fluorescence microscopy, the membranes were labeled with a 0.1 mol% of ATTO 647N-DOPE. **Figure 2a** shows that by tuning membrane charge and the salinity of the medium the interaction between membranes and the K_10_/D_10_ coacervates can be modulated. As previously shown for glycinin condensates,(48) and other complex coacervates(37), two wetting transitions can occur, from de-wetting (that is, no interaction), to partial wetting and complete wetting (complete spreading of the droplet over the membrane surface). Wetting of the membrane by biomolecular condensates imply mutual remodeling, since the condensate spreads on the membrane when the affinity is increased, and the membrane gets locally reshaped.(36, 37, 39, 50)

**Figure 2.**
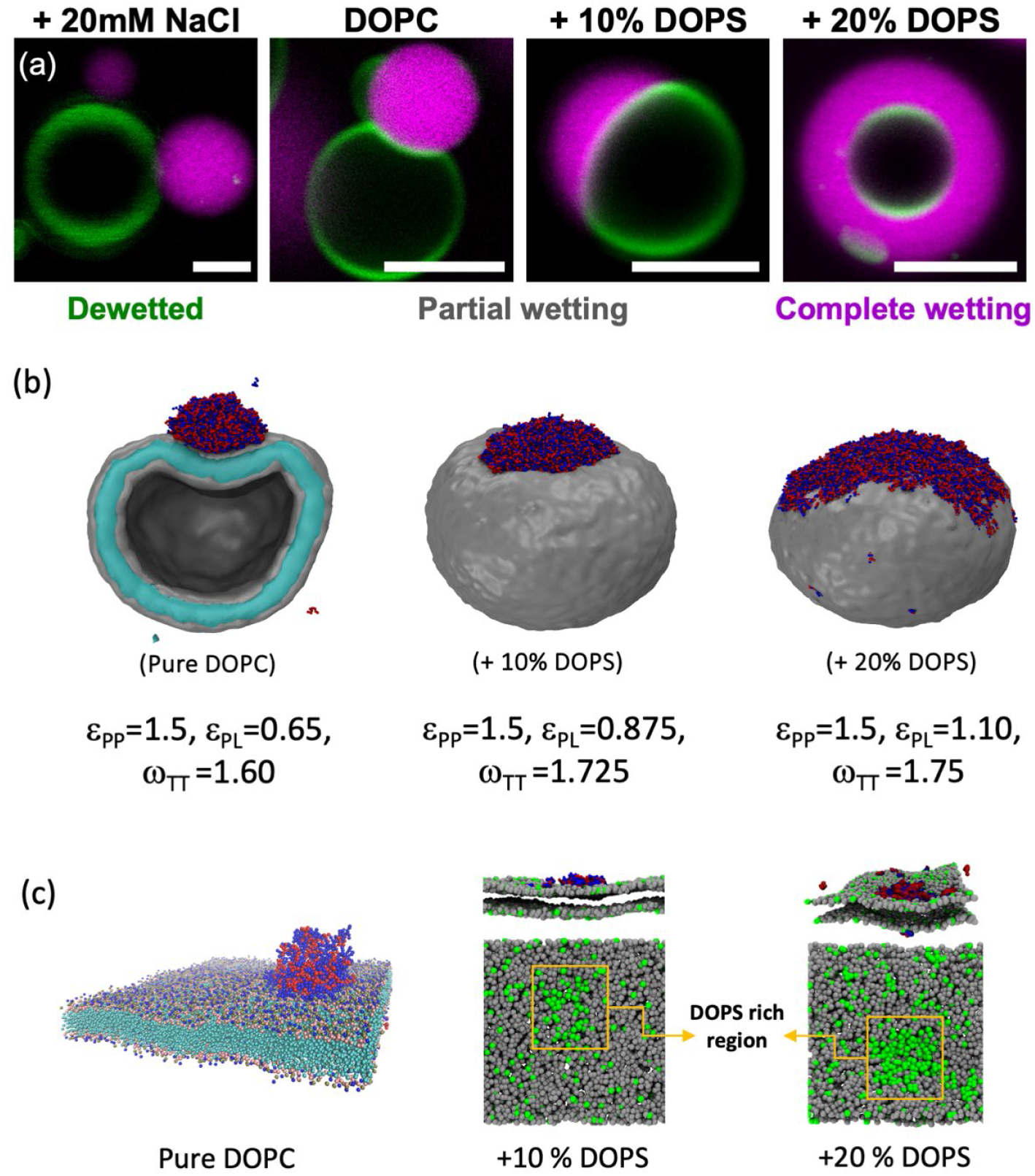
Wetting and lipid redistribution in membrane-coacervate interactions. (a) Confocal fluorescence images of DOPC or DOPC:DOPS GUVs labeled with 0.1 mol% ATTO 647N-DOPE (green) in contact with the K_10_/D_10_ coacervates (magenta) at the indicated membrane compositions. NaCl was added to pure DOPC vesicles when indicated (first image). In all cases, the final concentrations of KCl and MgCl_2_ were 15 mM and 0.5 mM, respectively. Scale bars: 5 µm. (b) Snapshots from the generic model molecular dynamics simulation at fixed value of ε_PP_ = 1.5 and different combinations of ε_PL_ and ω_TT_ which correspond to the systems shown in panel a as indicated above the snapshots. (c) K_10_/D_10_-coacervate in contact with bilayer patches of three different composition: Wetting, negative curvature generation, and local lipid demixing upon the adsorption of a K_10_/D_10_-coacervate on a DOPC:DOPS=9:1 and DOPC:DOPS=8:2 bilayer patches. Approximately 50 % of the charged lipids (green) in the leaflet facing the coacervate accumulate in the contact area.

In the generic CG model simulations using the generic Cooke lipid model, we explored the parameter space by keeping the interaction between peptides fixed at *ε*_*PP*_ = 1.5 and by tuning the peptide-lipid (*ε*_*PL*_) and lipid-lipid interaction range (*ω*_*TT*_). The objective was to optimize these interaction parameters to match the geometric factor, i.e. the intrinsic contact angle, with experimentally measured values. Once refined, these parameters enable us to leverage the predictive power of the simulations to resolve the structural rearrangements in the bilayer and changes in lipid dynamics upon wetting by the coacervates.

The optimized parameters are shown below **Figure 2b**. The three systems correspond to pure DOPC, 10% DOPS, and 20% DOPS compositions, respectively. In addition, we simulated another system that corresponds to 5% DOPS with *ε*_*PL*_ = 0.7625 and *ω*_*TT*_ = 1.70 (not shown in **Figure 2b**). The systems exhibit different extent of wetting. From MARTINI-3 simulations (**Figure 2c**) we studied three systems, namely pure DOPC, 10% DOPS, and 20% DOPS to probe the spatial distribution of lipids upon wetting by the coacervate. We observed that the coacervate induced local demixing around the area of contact and generated a negative curvature (from lumen’s perspective) on the 10% DOPS and 20% DOPS membrane (**Figure 2c** *middle and right panes*). We further quantified that almost 50% of the anionic lipids (DOPS) in the bilayer leaflet facing the coacervate are in contact with it, indicating a strong sorting and local demixing of the membrane. This local demixing and remodeling of the membrane could be crucial for processes like signaling and information transduction across the bilayer.(8, 87) Moreover, it has been shown that the electric potential gradient generated by the condensate formation can influence the membrane potential, offering additional means for membrane regulation.(88)

### 2. Quantifying the membrane-coacervate interaction at the micro-and nanoscales

At the micron scale, the vesicle membrane displays a “kink” at the contact line between the membrane, the coacervate phase, and the external solution, as illustrated in the partial wetting examples shown in **Figures 2a** and **3a**. This kink cannot persist at the nanometer scale, as it would imply infinite bending energy. Contact angles measured from microscopy reflect the overall geometry of the vesicle-condensate system and depend on their areas and volumes. Under identical conditions, vesicle-condensate pairs in the same sample may exhibit different microscopic contact angles. However, at the nanometer scale, the membrane must exhibit a smooth curvature (see **Figure 3b**), including an intrinsic contact angle with the condensate interface, as shown previously.(70, 89) The smooth curvature of the membrane has been also experimentally confirmed using super-resolution microscopy(69) and is also evident in the simulation snapshots, see **Figure 3c**.

**Figure 3.**
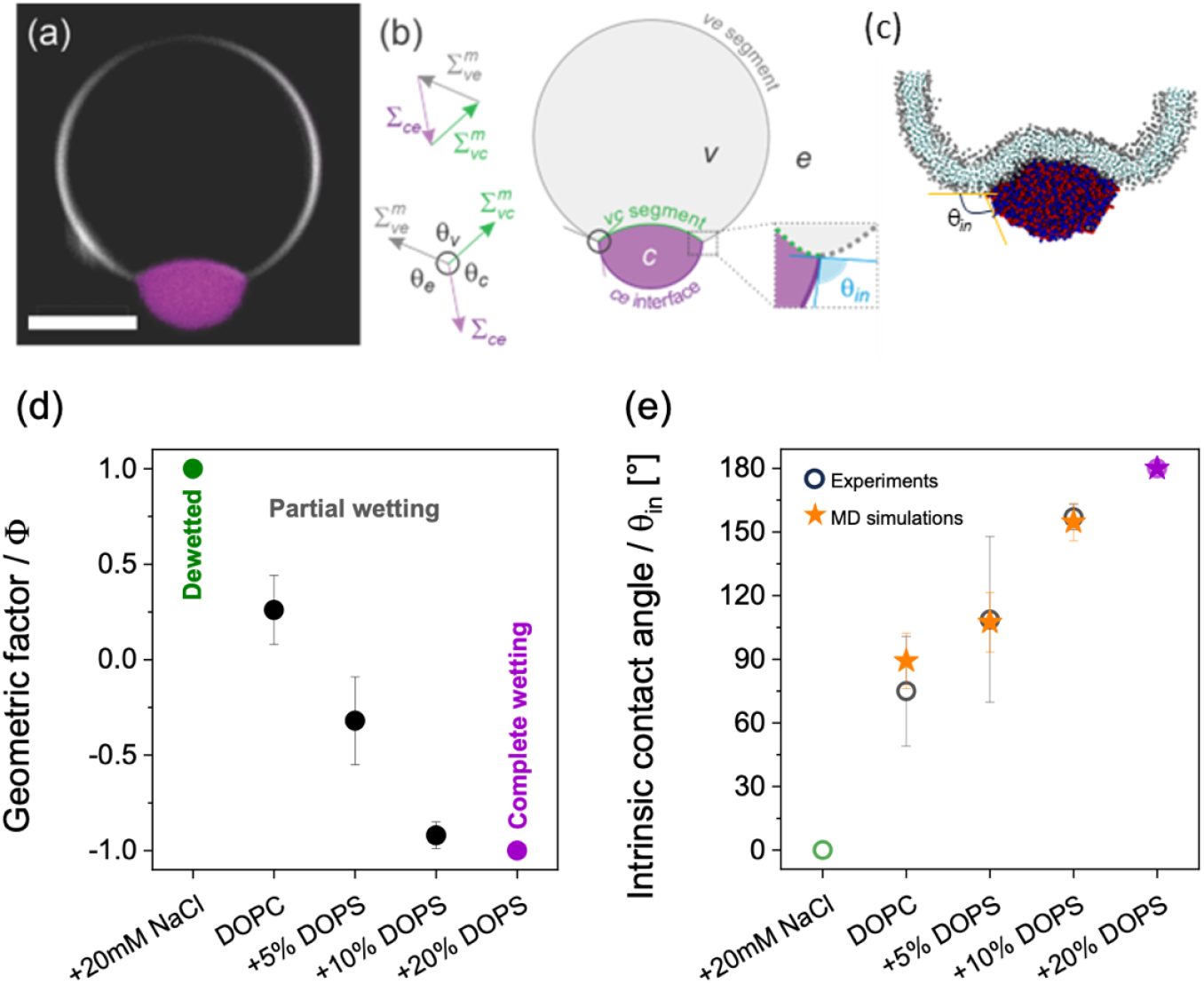
Wetting morphologies and intrinsic contact angles in vesicle-coacervate systems. (a) DOPC GUV in contact with a K_10_/D_10_ coacervate. Scale bar: 5 µm. (b) For partial wetting morphologies, the contact line between the condensate bare surface, ce, and the vesicle partitions the membrane into the vc and ve segments (green and gray, respectively), with the microscopic contact angles θ_e_ + θ_c_ + θ_v_ = 360^°^. The droplet interfacial tension Σ_ce_ and the mechanical tensions 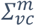 and 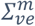 of the two membrane segments are balanced and form the sides of a triangle. The contact angles can be determined from the confocal cross-sections, and the geometric factor, Φ, can be calculated according to Eq. 1. (c) The intrinsic contact angle, θ_in_ as assessed from a simulation snapshot. (d) Geometric factor, Φ, experimentally estimated for vesicles of different membrane and milieu compositions. The system undergoes two wetting transitions, from dewetted ( Φ = 1) to partial wetting (™1 < Φ < 1) and complete wetting ( Φ = ™1) as a function of membrane surface charge and solution salinity. (e) Intrinsic contact angle θ_in_ from the confocal microscopy images (hollow circles), for the systems shown in d and from simulation snapshots (yellow stars) with the generic CG model and Cooke lipids. As the model does not include the effect of ions, we omit the leftmost point in simulations which represents complete dewetting.

The intrinsic contact angle is a material property, unlike the microscopic contact angles, and is uniform for all condensate-vesicle pairs in the sample under fixed conditions.(37, 44, 67) At the three-phase contact line, the force balance between the condensate interfacial tension (Σ_*ce*_) and the tensions of the two vesicle membrane segments (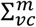and 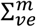) forms a triangle (analogous to the Neumann triangle),(68) see **Figure 3b**. We will refer to these microscopically measured contact angles as “apparent” because they do not persist at the nanoscale. By considering the force balance and that the membrane has different affinities for the condensate and external phases, the intrinsic contact angle can be estimated from measurements of the apparent microscopic contact angles, *θ*_*v*_, *θ*_*e*_ and *θ*_*c*_.(39, 70, 89) This relationship is given by **Eq. (1)**, where we introduce the geometric factor Φ = cos *θ*_*in*_, which reflects the membrane affinity for the two phases.(39, 70)

**Figures 3d** and **3e** show the quantification of the geometric factor and the intrinsic contact angle for K_10_/D_10_ coacervates in contact with DOPC and DOPC:DOPS vesicles. For complete dewetting, Φ = 1 and *θ*_*in*_ = 0°, for partial wetting −1 < Φ < 1 and 0° < *θ*_*in*_ < 180°, and for complete wetting Φ = −1 and *θ*_*in*_ = 180°. From the generic CG model simulations, we calculate the values of *θ*_*in*_ from several simulation snapshots across the trajectory and plot them in **Figure 3e** (yellow stars). With carefully chosen peptide-lipid and lipid-lipid interaction parameters, the obtained values (with error bars) corroborate well the experimental values (hollow circles in **Figure 3e**). To measure the angle, we took advantage of image processing tools namely, the Canny edge detection algorithm;^65^ see **Supporting Information** (SI, **Figure S3**) for details.

As we will demonstrate in the following sections, the strong agreement between the experimental results and the simulations in describing condensate-membrane interactions not only reinforces the observed experimental findings but also provides a complementary, molecular-level perspective on these interfaces.

### 3. Membrane dynamics at the contact region

Previous studies have reported that lipid dynamics can slow down upon contact with a condensate.(37, 87, 90) To determine whether this effect can also occur for the system explored here, we combined experimental and simulation data to compare lipid diffusion in the membrane region wetted by the coacervate with that in the bare membrane. Experimentally, we used fluorescence recovery after photobleaching (FRAP), as shown in **Figure 4a**. The results indicate that lipid diffusion in the wetted region is approximately 1.7 times slower than in the bare membrane. A similar reduction in diffusion is observed in vesicles composed of pure DOPC, as shown in the SI, **Figure S1.a**).

**Figure 4.**
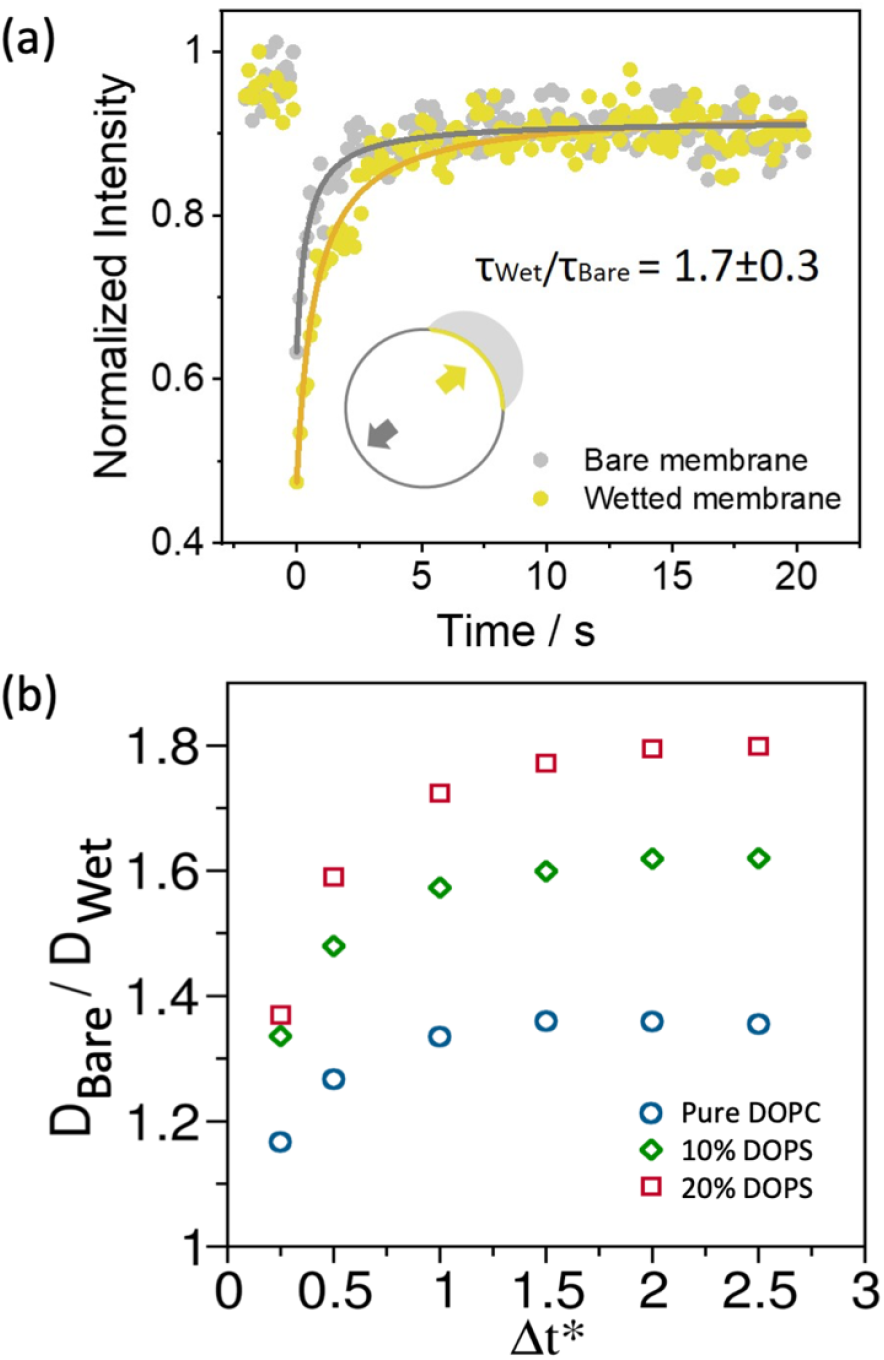
Lipid diffusion in the membrane region wetted by the coacervate is slower than in the bare membrane. (a) Fluorescence recovery after photobleaching (FRAP) of the fluorescent probe ATTO 647N-DOPE in DOPC:DOPS (9:1) membrane segments wetted by the coacervate (yellow) and the bare membrane (gray). Curves were fitted to the function: y=(I_0_+I_max_(x/τ_1/2_))/(1+x/τ_1/2_), where I_0_ is the initial intensity, I_max_ is the maximal intensity and τ_1/2_ is the halftime of recovery. (b) Ratio of lipid diffusion constants between the wetted and the bare membrane regions as obtained from the generic model CG simulations for the three systems (see **Figure 2b**).

From the generic CG model simulations, we indirectly estimate the ratio of the lipid diffusion constants in the wetted and the bare regions, *D*_*Wet*_ and *D*_*Bare*_, respectively. A direct calculation of the diffusion constants was not feasible because the residence time of the lipids at the contact region is too short to capture the long timescale behavior of the mean squared displacement. The velocity autocorrelation formalism cannot be applied as time in these CG models has no direct connection with real time.

To estimate the diffusion ratio, we measured the displacement of the lipid headgroups over a time interval Δ*t*^∗^ for lipids in contact with the coacervate as well as those far from it. Δ*tt*^∗^ was varied from 0.25 to 3.0. For each value of Δ*t*^∗^, the ratio of the squared total displacements provides an estimate of the ratio *D*_*Baree*_/*D*_*Wet*_. As shown in **Figure 4b**, this ratio saturates near 1.8 for the 20% DOPS system. This is in close agreement with the FRAP results, further validating the model parameters.

Since the reduced lipid mobility at the condensate-membrane interface may be linked to increased lipids ordering, as previously suggested,(37, 50) in the next section we explore that possibility.

### 4. Membrane packing and lipid ordering at the contact region

To assess whether lipid packing increases locally in our system, we labeled the membranes with a 0.5 mol% of LAURDAN and placed them in contact with unlabeled coacervates. LAURDAN is an environment-sensitive dye whose spectral properties respond to changes in lipid packing and hydration.(91) To evaluate its spectral shifts, we employed hyperspectral imaging combined with phasor analysis.(71)

Hyperspectral imaging allows us to capture spectral information at every pixel. Spectral phasor analysis, which involves applying a Fourier transform to the hyperspectral data (see methods), provides a straightforward visualization and quantification of LAURDAN spectral changes. These methods have been previously employed to characterize membrane-condensate interactions(37) and details on their application to membrane systems can be found in Ref.(71).

**Figure 5a** shows an example of a GUV composed of DOPC:DOPS (9:1) labeled with LAURDAN and in contact with three unlabeled K_10_/D_10_ coacervate droplets. The corresponding spectral phasor plot shown in **Figure 5b** reveals distinct contributions from the wetted and bare membrane regions, indicating that LAURDAN is sensing two different local environments. Pixel distribution histograms for these membrane segments, obtained using the two-cursor analysis (see Methods), are shown in **Figure 5c**, with their centers of mass plotted in **Figure 5d**. These results clearly indicate that the membrane packing is enhanced in the membrane segment interacting with the coacervate compared to the bare membrane (SI, **Figure S1.b-c** for membranes made of pure DOPC).

**Figure 5.**
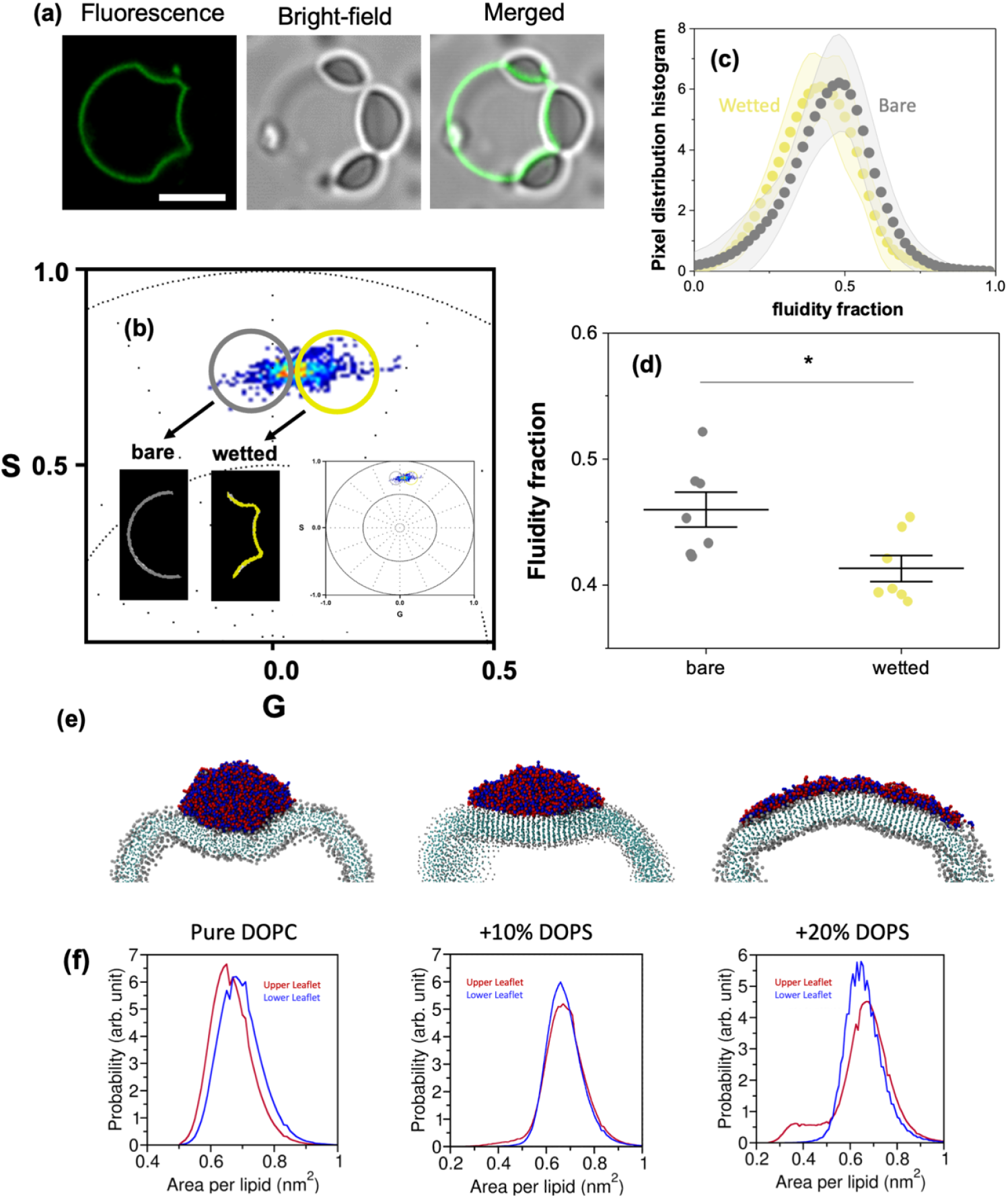
Lipid packing and ordering at the coacervate-membrane interface (a) GUVs composed of DOPC:DOPS (9:1) were labeled with 0.5 mol% LAURDAN and placed in contact with unlabeled K_10_/D_10_ coacervates. The images show an example of a GUV (green) wetted by three condensate droplets visible in bright field. Scale bar: 5 µm. (b) Spectral phasor plot for the vesicle displayed in (a). Using color cursor selection, the wetted (yellow) and bare (gray) membrane segments can be distinguished. The wetted membrane segment exhibits a blue shift, indicative of increased lipid packing. (c) Pixel distribution histograms for the wetted and bare membrane segments, presented as mean ± SD (n=8). (d) Center of mass of histograms shown in (c). The wetted membrane segments display a reduced fluidity fraction (higher packing) compared to the bare segments. Individual data points are shown as circles, and the lines correspond to mean ± SD. Statistical significance was determined using ANOVA and Tukey post-test (p<0.05). (e) Cross-sectional views of coacervates adsorbed on vesicles for three simulated systems showing orientational ordering of lipid tails at the coacervate-membrane contact region. (f) Distributions of the area per lipid (APL) for the pure DOPC, 10% DOPS, and 20% DOPS systems. The leaflet in contact with the coacervate, termed the upper leaflet (red), shows a decrease in the APL values, i.e. increased packing.

When comparing the spectral phasors of vesicles composed of DOPC or DOPC:DOPS (9:1) in the absence of coacervates, we observed that incorporating 10 mol% of DOPS results in a decrease in lipid packing (**SI, Figure S2**). Considering that DOPS is recruited to the membrane-coacervate interface (as shown in simulations, see **Figure 2c**), and that it has a larger (and charged) headgroup than DOPC, one might expect that electrostatic repulsion would reduce lipid packing in the contact region. However, we observed the opposite effect: lipid packing increased in the wetted membrane segment, which agrees with the reduced lipid dynamics observed in **Figure 4**. These results further support previous findings,(48) suggesting a general consequence of coacervate-membrane interaction.

In the Cooke lipid simulations, we observed orientational ordering of the lipid tails at the coacervate-membrane interface for all systems except for the one corresponding to pure DOPC liposome (**Figure 5e**). This ordering translates into tighter lipid packing in the contact region and provides enthalpic stabilization through increased interactions with the coacervate. The observed local reorganization of membrane lipids associated with increased order is consistent with the membrane dehydration(48) and transmembrane coupling reported previously.(87)

To further investigate lipid packing, we analyzed the area per lipid (APL) distribution using MARTINI-3 simulations for both the upper leaflet (in contact with the coacervate) and the lower leaflet (**Figure 5f**). For the pure DOPC system, we find a slightly left shifted APL distribution for the upper leaflet (**Figure 5f**, *left panel*). For the 10% DOPS bilayer, the APL distribution for the upper leaflet shows an increase in lipid population with lower values of APL below 0.5 nm^2^ (**Figure 5f**, *middle panel*). Interestingly, in the 20% DOPS system, the APL distribution for the upper leaflet includes a significant population with values below 0.5 nm^2^, indicative of a compact packing (**Figure 5f**, *right panel*). However, unlike the Cooke lipid model, the MARTINI bilayers do not exhibit significant lipid tail ordering.

While optical microscopy provides only limited insight into phase separation and the different emerging functions,(21) the integration of advanced microscopy techniques with molecular dynamics simulations, as demonstrated here, offers a powerful approach to understanding the interactions between membrane-bound and membraneless moieties (as represented by the vesicles and coacervates here) at the nano-and molecular scale. A similar experimental strategy, employing fluorescence lifetime microscopy (FLIM) has shown that the micro-polarity of condensates is determinant for their structural organization such as the formation of multilayered condensates in organelles like the nucleolus.(92) In addition, measurements of fluorescence anisotropy can reveal molecular interactions within condensate during LLPS providing understanding in their material properties.(93)

In this manner, the application of quantitative fluorescence microscopy techniques offers deeper insight into the diverse processes governing and involved in LLPS and the interactions with various organelles. The combination of these experimental approaches with molecular dynamics simulations enables a more comprehensive picture, extending resolution to the molecular scale.

## CONCLUSION

In this work, we have investigated the intricate interplay between coacervates and lipid membranes, an interaction fundamental to many cellular processes and the development of synthetic systems. By combining experimental observations of GUVs and K_10_/D_10_ coacervates with molecular dynamics simulations, we constructed an effective framework for elucidating the underlying mechanisms governing this complex phenomenon.

First, we successfully parameterized the coarse-grained model to accurately reproduce the fluid-elastic parameters observed in experiments, a critical step in establishing a reliable computational representation of the system. This allowed us to describe the wetting behavior of K_10_/D_10_ coacervates on membranes, characterized by the apparent contact angles and geometric factors at the microscale, as well as the intrinsic contact angle at the nanometric scale. When evaluating the membrane dynamic properties, our simulations further captured the experimentally observed mobility ratio between wetted and bare membrane regions, validating the approach of calibrating the coarse-grained model based on experimental contact angle parameters.

To elucidate the mechanism behind the reduced membrane fluidity at the coacervate interface, we performed hyperspectral imaging of LAURDAN combined with phasor analysis and demonstrated at the single-vesicle level that lipid packing increases in the wetted membrane segment compared to the bare membrane. Simulations provided molecular insight into this behavior, showing that ordering of lipids in the wetted region leads to a decrease in the area per lipid and demixing. Overall, our multiscale approach demonstrated that experiments and simulations not only yield consistent information, but also provide complementary perspectives, offering a richer view of these systems.

This work represents a significant advancement in our ability to quantitatively describe coacervate-membrane interactions. These interactions are not only important for membrane remodeling processes, but also can be key in signaling events, since both the lipid demixing and reduction of lipid dynamics at the contact region point to the formation of organized structures with distinct properties.

Our results contribute to a deeper understanding of the factors influencing coacervate-membrane interactions, paving the way for future studies exploring the impact of different coacervate compositions, membrane properties, and environmental conditions on these interactions. For example, it is of great interest to study the effect of cholesterol, which plays major roles in modulating the phase behaviors of cell membranes and the interaction between proteins and lipids.(94, 95) Indeed, a thorough analysis of interactions between protein condensates and raft-like lipid domains will help better understand the molecular mechanisms of cell signaling and its regulation.(8, 54, 59)For computational studies in this context, while a MARTINI description of cholesterol is available, a model in the framework of the Cooke lipids needs to be developed and calibrated carefully against experimental observations, which is worthwhile as the simulation of phase behaviors of multi-component lipid membrane is still computationally demanding with the MARTINI model. Another topic of interest concerns the relative importance of intrinsically disordered regions and structured domains to condensate/membrane interactions, since many proteins interacting with lipid membranes feature both disordered and structured regions.(96) Our present study provides a sound basis for future studies in this exciting, rapidly developing area by offering a robust framework integrating experimental and computational approaches for the description of the structural and dynamic properties at the interface. Along this line, further integrating with non-linear spectroscopic techniques and atomistic simulations that provide complementary level information about the structure and dynamics at such interface will be valuable.(47, 97, 98)

## ACKNOWLEDGEMENTS

A.M. acknowledges support from the Alexander von Humboldt foundation. Q.C. acknowledges the NSF grant, CHE-2154804. S.M. and Q.C. thanks the computing resources provided by the Shared Computing Cluster (SCC) at Boston University. R.D. acknowledges the ComeInCell network funded by the European Union’s Horizon Europe research and innovation program under the Marie Skłodowska-Curie grant agreement No. 101168939.

