## Supporting Information for "Integrating experiments and simulations to unravel coacervate-membrane Interactions: Insights into de-mixing and morphology modulation"

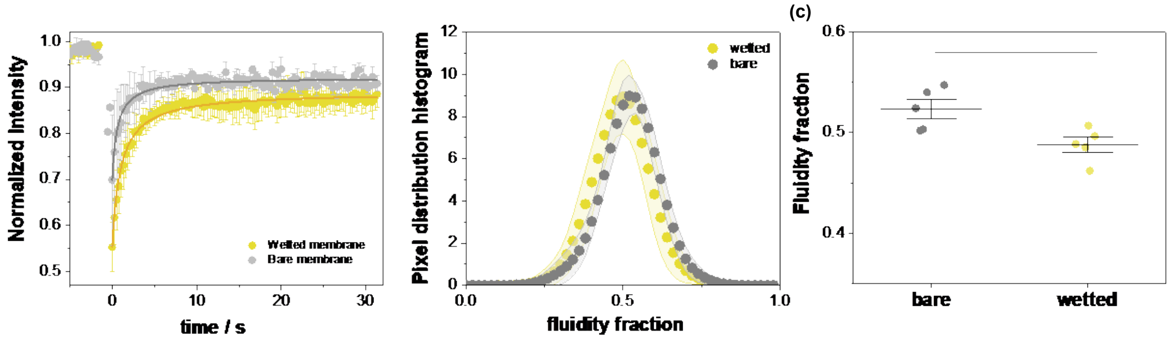


**Figure S1: Effects on diffusion coefficient and membrane packing for pure DOPC GUVs in contact with $K_{10}/D_{10}$ coacervates at 15 mM KCl and 0.5mM $MgCl_2$. (a) FRAP of the ATTO 647N-DOPE dye on membrane segments wetted by the coacervate (yellow) and the bare membrane (gray). Data are shown as mean±SD, n=10. (b) Pixel distribution histograms for the membrane wetted and bare segments. The histograms are shown as mean±SD (n=5). (c) Center of mass of the histograms shown in (c). The wetted membrane segments display a reduced fluidity fraction (higher packing) compared to the bare segments. Individual data points are shown as circles and the lines correspond to mean±SD. The differences are significant, $p<0.05$, ANOVA and Tukey post-test analysis.**

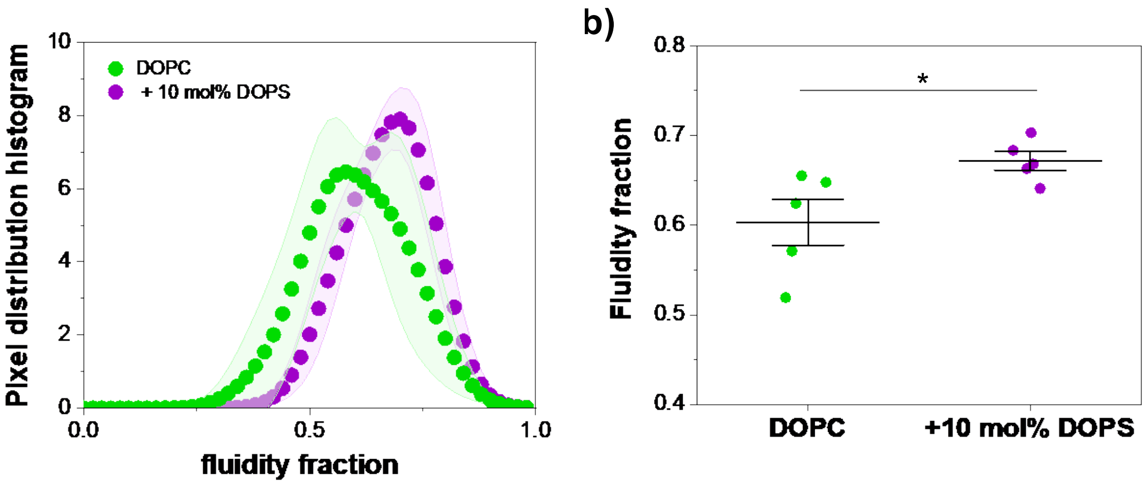


**Figure S2: Membrane packing measured for DOPC and DOPC:DOPS 9:1 membranes labeled with 0.5 mol% LAURDAN (in absence of coacervates). (a) Pixel distribution histograms for the DOPC and DOPC:DOPS 9:1 vesicles. The histograms are shown as mean±SD (n=5). (b) Center of mass of the histograms shown in (a). Adding DOPS increases the fluidity fraction (lower packing) compared to the pure DOPC membranes. Individual data points are shown as circles and the lines correspond to mean±SD. The differences are significant, p<0.05, ANOVA and Tukey post-test analysis.**

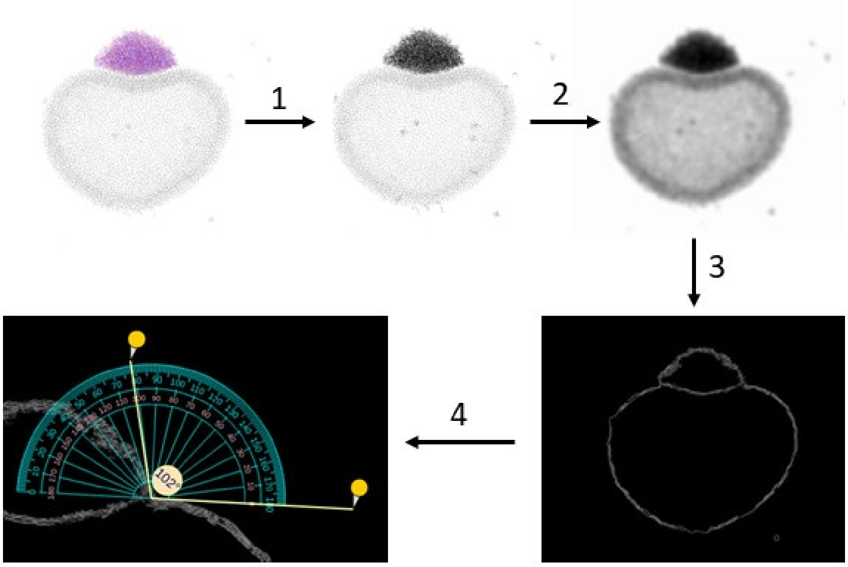

**Figure S3. Contact angle determination from simulation snapshot: First we find a suitable two-dimensional projection using Visual Molecular Dynamics (VMD) software and convert the colored image into a greyscale one (Step-1). In Step-2, a gaussian blur filter is applied along with gamma correction to produce the input for the edge detection. In Step-3, Canny edge detection algorithm is applied to the output of Step-2. Finally, in Step-4, a web-tool named 'online protractor' ([https://www.ginifab.com/feeds/angle_measurement/](https://www.ginifab.com/feeds/angle_measurement/)) is used to measure the intrinsic contact angle.**
